# Genetic variations (eQTLs) in muscle transcriptome and mitochondrial genes, and trans-eQTL molecular pathways in feed efficiency from Danish breeding pigs

**DOI:** 10.1101/2020.04.17.047027

**Authors:** Victor AO. Carmelo, Haja N. Kadarmideen

**Author notes:** **Correspondence:** Haja N. Kadarmideen.

## Abstract

Feed efficiency (FE) is a key trait in pig production, as it has both high economic and environmental impact. FE is a challenging phenotype to study, as it is complex and affected by many factors, such as metabolism, growth and activity level. Furthermore, testing for FE is expensive, as it requires costly equipment to measure feed intake of individual animals, making FE biomarkers valuable. Therefore, there has been a desire to find single nucleotide polymorphisms (SNPs) as biomarkers, to assist with improved selection and improve our biological understanding of FE. We have done a cis- and trans-eQTL (expressed quantitative trait loci) analysis, in a population of Danbred Durocs (N=11) and Danbred Landrace (N=27) using both a linear and Anova model. We used bootstrapping and enrichment analysis to validate and analyze our detected eQTLs. We identified 15 eQTLs with FDR < 0.01, affecting several genes found in previous studies of commercial pig breeds. Examples include IFI6, PRPF39, TMEM222, CSRNP1,PARK7 and MFF. The bootstrapping results showed statistically significant enrichment of eQTLs with p-value < 0.01 (p-value < 2.2×0^-16^) in both cis and trans-eQTLs. Based on this, enrichment analysis of top trans-eQTLs revealed high enrichment for gene categories and gene ontologies associated with genomic context and expression regulation. This includes transcription factors (p-value=1.0×10^-13^), DNA-binding (GO:0003677, p-value=8.9×10^-14^), DNA-binding transcription factor activity (GO:0003700,) nucleus gene (GO:0005634, p-value<2.2×10^-16^), positive regulation of expression (GO:0010628), negative regulation of expression (GO:0010629, p-value<2.2×10^-16^). These results would be useful for future genome assisted breeding of pigs to improve FE, and in the improved understanding of the functional mechanism of trans-eQTLs.

## Introduction

The biological background of complex traits is expressed through molecular processes triggered by a combination of genetics, epigenetics and the environment. While ample genetic markers have been identified for complex traits, the understanding of the functional effect of identified genetic markers is challenging to identify(1). Almost per definition, complex traits are controlled by multiple genetic factors (2–4), thus further complicating the biological control mechanisms. One way of tackling this issue, is to look at direct causal links between genetics and gene expression, thus identifying a direct effect of genetic variation. This allows for a straightforward interpretation of the effect of genetic variation based on pathway and functional knowledge of implicated genes. This can be done through the identification of expressed quantitative loci (eQTL), mapping genetic variants that influence gene expression patterns of genes in various tissues, originally termed as systems genetics (5, 6). The usage of both the genetic and the transcriptomic information, combined with pathway and phenotype data can be a powerful way of identifying biomarkers for traits of interest. There are however, several challenges with eQTL analysis. Firstly, if one wanted to map all possible SNP-gene pairs in a modern data set, which typically has thousands of expressed genes and at the minimum several tens of thousands of SNP, the total amount of tests will be at least in the order of 10^8^. This can pose computational challenges, but even worse, multiple testing problems. This is especially relevant as a cursory search of the Gene Expression Omnibus database (https://www.ncbi.nlm.nih.gov/geo/) for RNA-seq studies reveals most studies having less than 100 samples. Therefore, it is important to have strategies for these issues when doing eQTL analysis. Example strategies used for filtering the expression data in previous studies include: filtering by estimated heritability of gene expression (7–9) or using only a limited set of genes(10), and there are many more possibilities.

Feed efficiency (FE) has been known for decades to be an important complex trait in pig breeding. Cost of feed is the largest economic burdens in commercial pig production (11, 12), and lower feed consumption leads to more environmentally friendly production. The two main metrics for feed efficiency are residual feed intake (RFI) (13) and feed conversion rate (FCR), which isthe ratio between feed consumed and growth), with the latter being the most used in pig production. Selective breeding has improved FCR in pigs, but this has not led to direct gains in knowledge of the biological drivers of FE in pigs. Even with many studies being done on the subject, the genetic and biological background of FE in pigs is still not well understood (14). The cost and difficulty of measuring FE likely plays into this, as it cannot be easily measured without expensive equipment and setup, unlike meat quality or litter size. One concrete example of the usage of FCR is in the Danish pig production where, FCR is improved through a centralized breeding program were potential breeding sires are tested for efficiency via accurate calculations based on measured feed intake and growth.

Muscle is the most important tissue in pig production in regards to economic value. Muscle plays a large role in energy metabolism and energy storage (15–17). As such, there have been multiple studies on the muscle transcriptome in a FE context (11, 18–20). In comparison, while there are several eQTL studies performed in pig muscle (7, 10, 21, 22), there are none based on FE traits. A connection between FE and mitochondria in muscle has been reported several times, in several species in the literature (11, 19, 20, 23–25). In general, it is reported that higher mitochondrial activity is related to increased FE. Given the evidence for mitochondrial effects, identifying genetic regulation of mitochondrial genes could assist in efforts to develop biomarkers for FE.

Trans-eQTLs are per definition distally located in relation to the genes they are affecting. This means that true trans-eQTLs should have a mechanism that mediates the correlation between expression and genetic variation. There has been evidence that trans-eQTLs can be mediated by local cis effects of the eQTL(26, 27). One proposed method for the mediation is through cis affected transcription factors (27). What is then the gene ontological nature of genes that are regulated by trans-eQTLs? We could hypothesize that the genes should be enriched for gene ontology categories that can interact with genomic context or regulate expression, as seen previously in genes near trans-eQTLs.

Here we aimed to perform both cis and trans-eQTL analysis on a previously identified set of FCR related differentially expressed genes (DEG) and mitochondrial genes (known to be involved in energy metabolism and nutrient utilization), in a pig population comprised of Danbred Duroc and Landrace purebred pigs. The two-breed analysis provides genetic variations that can aid in the detection of eQTLs (28), particularly as the Durocs were more heavily selected for FCR. By focusing on DEG and mitochondrial genes, we applied a targeted approach for underpinning systems genetics of our phenotype of interest (FE). To improve the statistical and computational analysis, we reduced genotype input space through linkage disequilibrium (LD) and loci variation filtering. Finally, we tested the hypothesis that genes that were associated with SNPs identified as trans-eQTLs would belong to pathways that could mediate such effects, including expression regulation and DNA binding.

## Material and Methods

### Sampling and Sequencing

The pigs in this study were the intersection between the pigs genotyped in Banerjee et al. (29) and Carmelo et al (30), resulting in a selection of 38 pigs. All data processing steps follow those two studies, unless otherwise stated. Of the 38 male uncastrated pigs included in this study,11 were purebred Danbred Duroc and 27 were purebred Danbred Landrace. The pigs were sent to the commercial breeding station at Bøgildgård, which is owned by the pig research Centre of the Danish Agriculture and Food Council (SEGES) at ~7kg. The pigs were regularly weighed, and feed intake was measured based on a single feeder setup in a test period from ~28kg of weight until ~100kg. The period of measurement was determined by each pig’s commercial viability.

### Data selection and filtering

All gene annotation and analysis was done using *Sus scrofa* annotation version 11.1.96 from Ensembl.

#### Gentoype Data and Filtering

The DNA isolation from collected blood and genotyping was performed by GeneSeek (Neogen company - https://www.neogen.com/uk/). The Genotyping was based on the GGP Porcine HD array (GeneSeek, Scotland, UK), which includes 68,516 SNPs on 18 autosomes and both sex chromosomes. The SNPs were mapped to the *Sus scrofa* genome version 11.1 using the NCBI Genome Remapping from the *Sus scrofa* genome version 10.2. This was done using default settings. To insure that we had a sufficient representation of genotypes for each SNP, we used a MAF (minor allele frequency) threshold of 0.3. This removes SNPs that would be underpowered for our eQTL analysis given our, or that cannot be related to expression changes due to lack of variation. It also has the advantage of reducing the overall testing space to a more conservatively sized set. This reduced the initial set of SNPs to a total of 27531. The next step performed was to remove groups of SNPs in high LD To do this, we used the *LD_blocks* function from the *WISH-R* R package(31), which was applied with an R^2^ of 0.9. This grouped SNPs linearly across chromosomes into blocks based on a minimum pairwise R^2^ value of 0.9 between all SNPs in a block. After this step, 19179 SNPs remained. The genotypes were coded as 0 (homozygote major), 1 (heterozygote) and 2 (homozygote minor) for the eQTL analysis.

#### Expression data, Gene selection and filtering

Muscle tissue samples were extracted from the psoas major muscle immediately post slaughter, and the samples were kept at −25 C in RNA later (Ambion, Austin, Texas). The data was sequenced on the BGISEQ platform using the PE100 (pair end, 100bp length) with RNA extraction and sequencing performed by BGI Genomics (https://www.bgi.com/global/). The mean number of total reads was 64.5 million with standard deviation of 7,4 × 10^5^. The mean number of uniquely mapped reads was 95.3% with a standard deviation of 0.33%. The reads were trimmed using Trimmomatic (32) version 0.39, with the default setting for paired end reads. Data QC was performed pre- and post-trimming using FastQC v0.11.9. Mapping was done with STAR aligner(33) version 2.7.1a, with default parameters and genome and annotation 11.1 version 96. Beside default parameters, the --quantMode GeneCounts setting was used for read quantification. Our main interest was to investigate genes that could be related to FCR. We therefore based our set of genes on the methods in Carmelo et. al (30). In brief, Differential Expression analysis (DEA) was performed using three different DE methods (Limma, EdgeR, Deseq2)(34–36) with FCR as the phenotype of interest. We then calculated the divergence between our observed p-value distribution for FCR and the uniform distribution for each method, enabling us to select a list of genes that are related FCR. This was motivated by the fact that we had a large overrepresentation of low p-values in the DEA, meaning the distribution was anti-conservative. This resulted in a set of 853 genes. As mitochondrial genes have been implicated in FE in muscle in both our previous study and in several studies in multiple species(11, 19, 20, 23–25), we also selected all genes with a mitochondrial gene ontology (gene ontology id GO:0005739) and included them in the analysis. All genes were filtered to have a minimum of 5 reads in at least 11 samples, as 11 was the size of the Duroc group. Testing revealed that genes with a single expression outlier could result in likely false positives. Therefore, all genes with a single gene with a Z-score above 3 were removed, corresponding to a single observation with normalized expression further than 3 standard deviations from the mean. This resulted in a final gene set of 1425 genes.

### eQTL Analysis

#### Calculation of eQTLs

All of the eQTL analysis was performed using R version 3.5.3. Gene expression was normalized using the *calcNormFactors* from the R package edgeR version 3.34.3. We performed eQTL analysis using the R package MatrixEQTL version 2.3(37). We added the following covariates in the model: RNA integrity values (RIN), breed, batch and age (days). Given that the samples were collected in slaughterhouse setting, it was necessary to include RIN in the model, but this should not be an issue if appropriately corrected for (38). As samples were collected on different days, it was necessary to correct for this using the batch effect. Breed and age have an effect on expression, as seen in our previous study (30) and thus must be corrected for. While the samples come from a selection of 28 different breeders in Denmark, there is still some relationship between some pigs, especially if they came from the same breeder. Therefore, a kinship matrix based on 4 generations of pedigree was added as the error covariance matrix instead of using the default identity matrix. The cis-distance was set to 10^6^ bp. The analysis was done using both the *modelANOVA* (Anova) and *modelLINEAR* (linear) options, giving both a factor based model and a linear model fit.

#### Statistical Significance

After the model was fit, based on the empirical p-value distribution, pathway enrichment analysis was performed on the top putative eQTLs based on the results from the trans-eQTL linear model. The linear model was chosen over the Anova as the empirical p-value distribution for the Anova had an overweight of low p-values, which means that we should avoid using the overall distribution of p-values for conclusions. In the linear version, we observed that the p-values were nearly uniform with a slight overweight of low p-values. To show the significance of this result, we performed bootstrapping by shuffling the genotype values of each SNP while maintaining the same expression values and covariates. We then calculated the number of random eQTLs with p-value < 0.01, for both the cis- and trans-eQTLs. Assuming the shuffled values are normally distributed, we calculated the probability of observing our empirical number of p-values < 0.01. We also saved the lowest, the 10th lowest and the 100th lowest observed p-value for both trans and cis bootstrapped eQTLs for each iteration. The bootstrapping procedure was done 500 times with both Anova and the linear model.

#### Orthonormalization

To visualize the expression and genotype values on the scale used by Matrix eQTL, we scaled and centered both the design matrix of the covariates, the expression and the genotypes. After this, we used the *mlr.orthogonalize* function from the MatchLinReg package version 0.7.0 to orthogonilize the expression values and genotypes of each relevant gene and SNP in relation to the covariates, respectively, using *normalize=True* as an option. This procedure was done mimicking the method reported in the Matrix eQTL(37).

#### QTL region and relation to FCR

To verify if our eQTLs were in know quantitative trait loci (QTL) regions, we first defined a region of 100kb upstream and downstream of each SNP to overlap with. The region size was conservatively defined based on reported haplotype block sizes in commercial pigs(39). We then checked if the SNP coordinate had any overlaps with FCR quantitative trait loci (QTL) from the Pig QTL database(40). We also did the same procedure with the the gene associated with each eQTL, except we did not extend the region beyond the gene boundaries.

#### Pathway Analysis

We hypothesized that, if trans-eQTLs are not false positives, they should be enriched for functional categories which could relevantly cause distal interactions, in comparison to the background. Therefore, to analyze our top trans-eQTLs, we calculated the number of additional empirically observed low p-values under 0.01, by subtracting the expected number of p-values < 0.01 given a uniform p-value distribution, from the observed number of p-values < 0.01. We then tested the enrichment of the genes in our top eQTL group for the following gene categories/ontologies: transcription factors(TF) (based on the AnimalTFDB 3.0 pig transcription factors (41)), DNA-binding (GO:0003677), DNA-binding transcription factor activity (GO:0003700) nucleus gene (GO:0005634), positive regulation of expression (GO:0010628), negative regulation of expression (GO:0010629) and membrane gene (GO:0016020). Each category was selected based on a biological hypothesis, with membrane gene serving as a kind of reference background category. All GO terms were retrieved using biomart 2.42.0 with annotation from *Sus scrofa* 11.1. 96

## Results

### eQTL analysis

In figure 1 we can see the overall p-value distribution for both the linear and the Anova eQTL analysis. The linear model is overall well behaved, with uniform p-values and a small increase of low p-values. In the Anova model, the spike of high p-values may be due to issues with model assumptions in some cases, but as there large number of eQTLs it is not practical to do model diagnostics on each eQTL. This does not mean individual Anova based eQTL cannot be valid, but we should be careful with drawing results based on the overall distribution. The cis-eQTLs have a more uneven overall distribution, but this likely due to the lower amount of tests combined with the histogram binning. Looking at the individual results using a threshold for FDR of 0.1, the only analysis that give any significant results was the Anova analysis, yielding 14 significant trans-eQTLs and 1 cis-eQTL. It should be noted, that in our linear analysis, due to the left skewing of the p-value distribution, all trans-eQTLs with p-value < 0.01 (N=301213) have and FDR value of 0.9 or better. This means that it is very likely that we have real trans-eQTLs, we just lack the power to identify them individually. Given this, and the results from the bootstrapping analysis (see below), we elected to show the top ten 10 eQTLs for each analysis, except the Anova trans, were we selected all with FDR < 0.1, which can be seen in table 1. In figure 2 we can see the visualization of the top 6 eQTLs in the linear trans model, ordered from lowest p-value (top left) to highest (bottom right). Given the low p-values reported, the visualization, especially of the lowest p-value, does not seem to support the results. The explanation is found in the way the Matrix eQTL implementation deals with covariates. In Matrix eQTL, all covariates, expression and genotypes are centered and scaled, and then the expression and genotype vectors are both orthogonized in relation to the covariate matrix. Only after this step is the linear relationship between expression and genotype calculated. In figure 3, we can see the same results as in figure 2, based on the transformed values. Here we can see a clear linear relationship between the transformed expression values and genotype values.

**Figure0 1.**
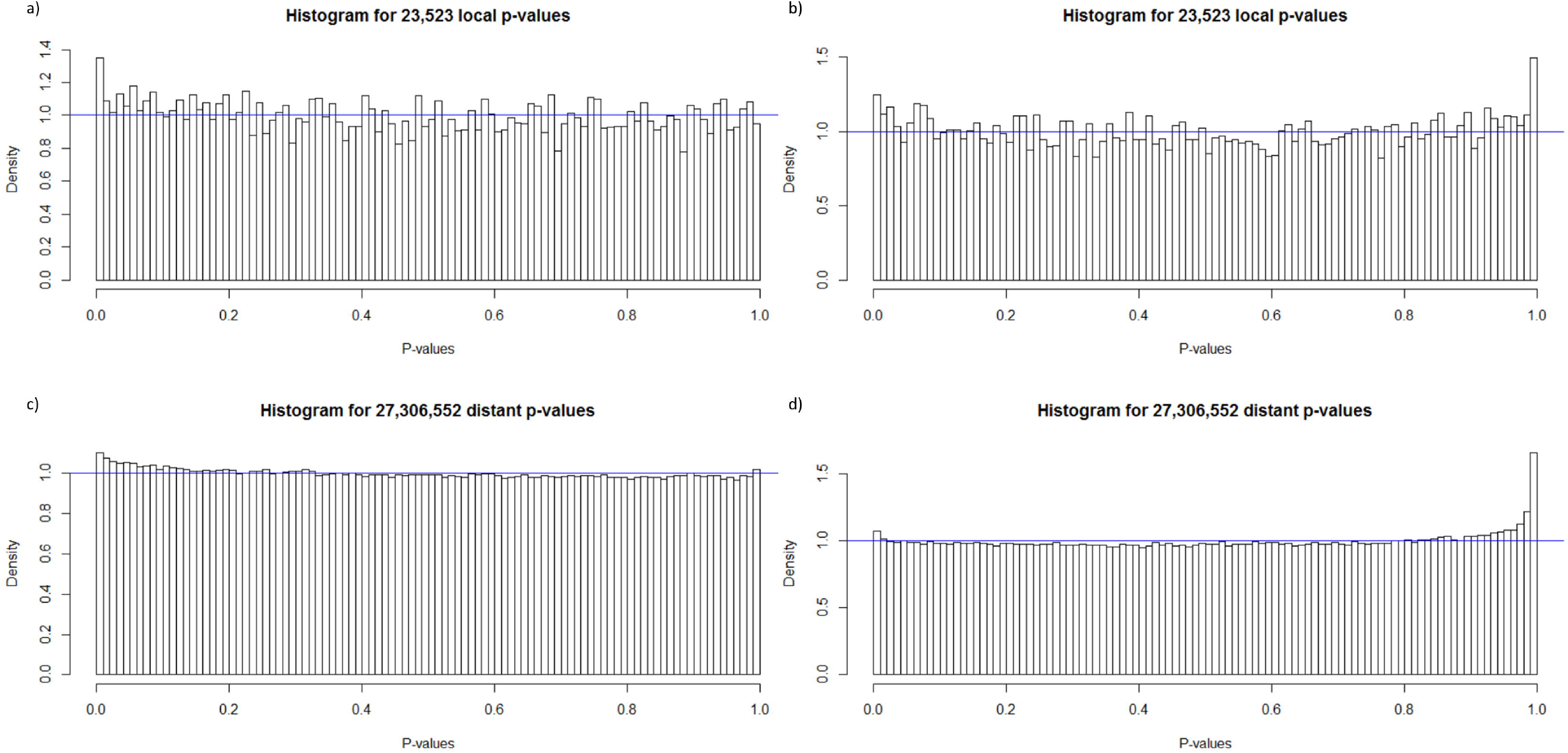
Histograms of the p-value distribution of all cis (a,b) and trans(c,d) eQTL pairs in the linear(a,b) and Anova(c,d) models. Based on the overall distribution, we see a slight anti-conservative trend in the linear p-values in both cis and trans-eQTLs.

**Table 1.** Overview over top cis and trans-eQTLs in all 4 four sub-analyses. ^1^Genes or SNPs in known FCR QTL regions. ^2^Snps with p-value < 0.05 for linear association with FCR

| Snp | Gene | P-value | FDR | Chr | Position | Analysis |
| --- | --- | --- | --- | --- | --- | --- |
| WU_10.2_7_1320670 | SLC20A2 | 2.36e-11 | 0.00064 | 7 | 1132634 | Anova trans |
| WU_10.2_7_115152142 | SLC20A2 | 1.21e-10 | 0.00096 | 7 | 108750676 | Anova trans |
| ASGA0040859 | SLC20A2 | 1.35e-10 | 0.00096 | 9 | 3330061 | Anova trans |
| WU_10.2_18_2886712 | SLC20A2 | 1.41e-10 | 0.00096 | 18 | 2875724 | Anova trans |
| ASGA0042452 | IFI6 | 7.68e-10 | 0.0042 | 9 | 31552392 | Anova trans |
| WU_10.2_14_148897646 | PRPF39 | 3.89e-09 | 0.018 | 14 | 137024504 | Anova trans |
| ALGA0109564 | DNAJB1 | 1.23e-08 | 0.048 | 15 | 68774343 | Anova trans |
| ALGA0053497 | TMEM222 | 1.40e-08 | 0.048 | 9 | 60196621 | Anova trans |
| WU_10.2_12_33709155 | GCAT | 2.01e-08 | 0.058 | 12 | 32811156 | Anova trans |
| ASGA0013363 | DLC1 | 2.11e-08 | 0.058 | 3 | 11038140 | Anova trans |
| ASGA0091484 | CSRNP1 | 2.55e-08 | 0.059 | 4 | 118914325 | Anova trans |
| ALGA0009614 | CSRNP1 | 2.58e-08 | 0.059 | 1 | 256190584 | Anova trans |
| WU_10.2_13_216907306 | POGZ | 3.21e-08 | 0.063 | 13 | 207030935 | Anova trans |
| ASGA0083137 | DLC1 | 3.22e-08 | 0.063 | 9 | 138505307 | Anova trans |
| ALGA0018160<sup>1</sup> | INTS7 | 5.61e-09 | 0.15 | 3 | 27307613 | Linear trans |
| MARC0081581<sup>1</sup> | INTS7 | 3.25e-08 | 0.31 | 3 | 27346598 | Linear trans |
| ALGA0015229<sup>2</sup> | ACOX3 | 3.39e-08 | 0.31 | 2 | 116633408 | Linear trans |
| ALGA0056299<sup>2</sup> | PARK7<sup>1</sup> | 5.21e-08 | 0.36 | 10 | 1400269 | Linear trans |
| ASGA0091638<sup>2</sup> | CPT1B | 8.19e-08 | 0.38 | 4 | 626787 | Linear trans |
| WU_10.2_12_3964486 | MFF<sup>1</sup> | 8.38e-08 | 0.38 | 12 | 4217210 | Linear trans |
| ALGA0115669<sup>2</sup> | PARK7<sup>1</sup> | 9.79e-08 | 0.38 | 10 | 1187360 | Linear trans |
| DRGA0015709<sup>1</sup> | COMTD1 | 1.22e-07 | 0.42 | 16 | 2090820 | Linear trans |
| ALGA0087901<sup>1</sup> | NSUN2 | 1.63e-07 | 0.45 | 15 | 129751572 | Linear trans |
| WU_10.2_7_740616 | Glycine N-phenylacetyltransferase | 1.64e-07 | 0.45 | 7 | 618465 | Linear trans |
| INRA0015708<sup>1</sup> | NES | 4.07e-06 | 0.096 | 4 | 94242001 | Anova cis |
| WU_10.2_15_134661069 | ABCB6 | 1.31e-05 | 0.11 | 15 | 121481724 | Anova cis |
| WU_10.2_6_27531636 | NUDT21 | 1.36e-05 | 0.11 | 6 | 30064483 | Anova cis |
| MARC0009689 | HDHD5 | 3.01e-05 | 0.18 | 5 | 69473204 | Anova cis |
| WU_10.2_15_150992806 | RAMP1 | 6.83e-05 | 0.32 | 15 | 136516507 | Anova cis |
| MARC0112128 | KCNMA1 | 0.00010 | 0.41 | 14 | 79922641 | Anova cis |
| WU_10.2_15_91334711 | MTX2 | 0.00022 | 0.55 | 15 | 81867240 | Anova cis |
| WU_10.2_2_4374745 | SYT12 | 0.00022 | 0.55 | 2 | 5445193 | Anova cis |
| ALGA0019808<sup>1</sup> | MEIS1 | 0.00026 | 0.55 | 3 | 76307479 | Anova cis |
| WU_10.2_3_18580686 | STX4 | 0.00027 | 0.55 | 3 | 18010007 | Anova cis |
| ASGA0054417 | MYO19 | 0.00018 | 0.63 | 12 | 38196853 | Linear cis |
| WU_10.2_X_128169493 | RBMX | 0.00018 | 0.63 | X | 112221790 | Linear cis |
| WU_10.2_12_39624033 | MYO19 | 0.00026 | 0.63 | 12 | 37981199 | Linear cis |
| WU_10.2_3_183721 | C7orf50 | 0.00031 | 0.63 | 3 | 335933 | Linear cis |
| ALGA0108896<sup>1</sup> | CRYM | 0.00035 | 0.63 | 3 | 24920076 | Linear cis |
| ALGA0061099 | MRPS31 | 0.00035 | 0.63 | 11 | 16202962 | Linear cis |
| ALGA0061107 | MRPS31 | 0.00035 | 0.63 | 11 | 16236530 | Linear cis |
| ASGA0030240 | NSUN4 | 0.00042 | 0.63 | 6 | 165835717 | Linear cis |
| WU_10.2_14_153092095 | ECHS1 | 0.00047 | 0.63 | 14 | 141129811 | Linear cis |
| WU_10.2_14_153836231 | ECHS1 | 0.00047 | 0.63 | 14 | 141357898 | Linear cis |

**Figure0 2.**
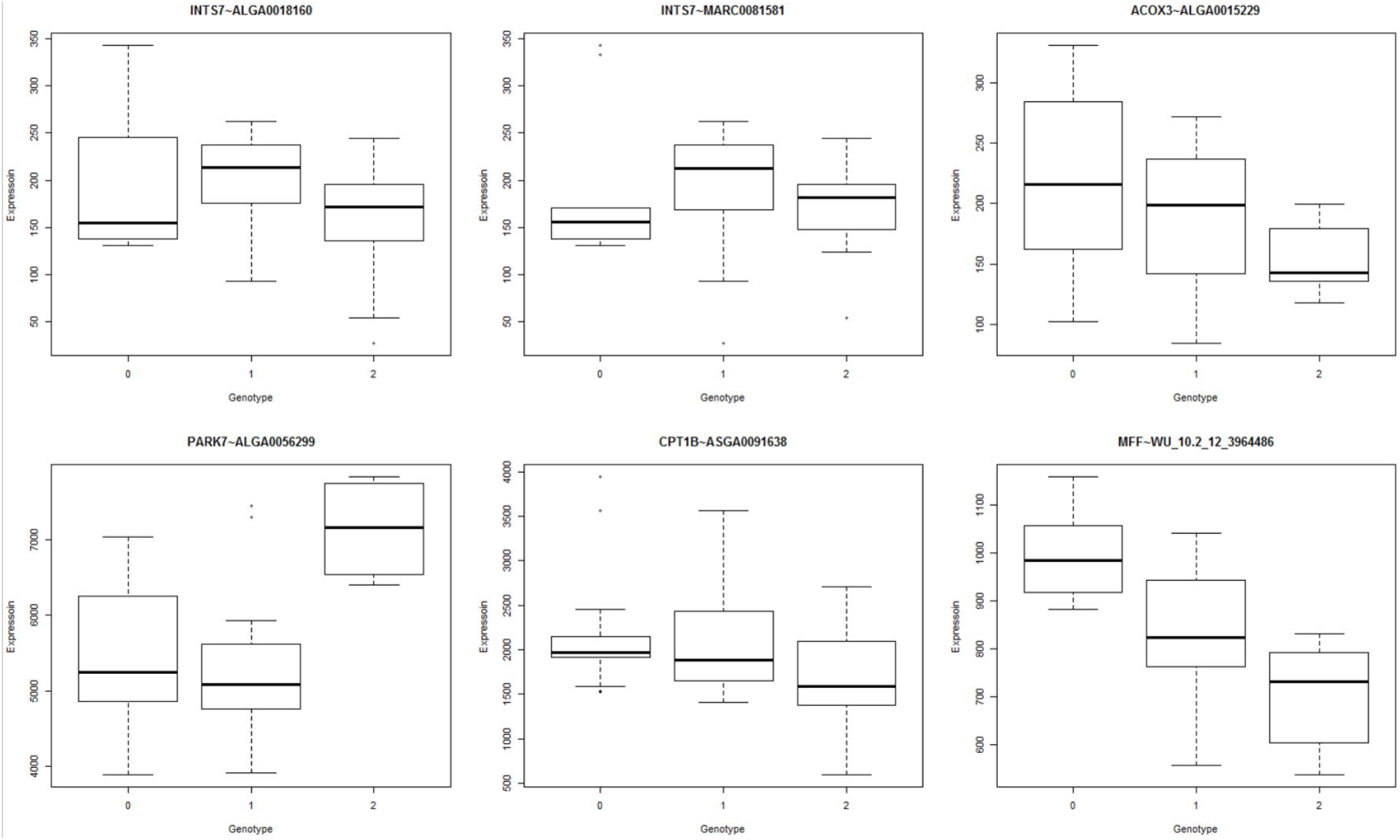
Boxplot of the top 6 trans-eQTLs from linear analysis. Comparing with the summary from table 1, it seems unexpected that the top left boxplot is of the most significant eQTL. Overall, the 3^rd^ and the 6^th^ ranked eQTLs look visually more appealing. This because the genotype here are the raw values, and the expression values are normalized but without taking covariates into consideration.

**Figure0 3.**
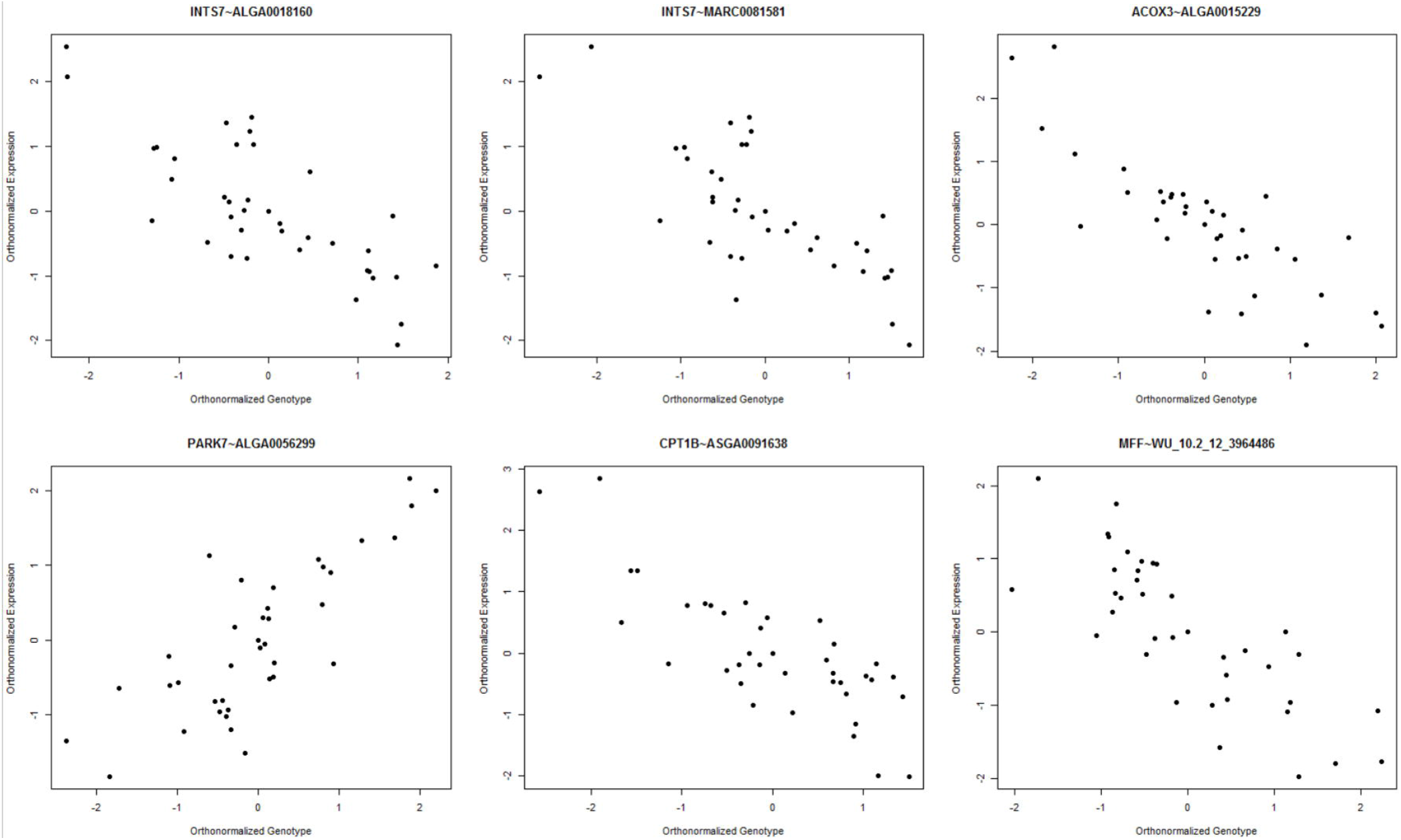
Scatter-plot of the orthonormalized expression and genotype values for the top 6 trans-eQTLs in the linear analysis. The linear relationship is quite clear on the transformed values, in comparison to the boxplots of the untransformed values.

#### Bootstrapping

Bootstrapping is a useful tool when dealing with complex data, allowing us to get estimates of the likelihood of our observations without explicit probability calculations. Here, we wanted to show that our spike in low p-values in the linear analysis was statistically unlikely to happen by chance. Based on 500 bootstraps, we estimated the mean and the variance of the number of p-values < 0.01, and compared this with our observed number. Assuming normally distributed counts, which is a fair assumption given our individual bootstrapping sample size and the scale, the probability of our observed value is essentially 0, which is also visualized in figure 4. In table 2 we can see the comparison of the 1^st^, 10^th^ and 100^th^ p-values in our bootstrapped data with our empirical data. Overall, the real data was significantly more left skewed as we go down in rank to the 100^th^ p-value. This indicates that the real data has lower bound on significance, but overall the results are not achievable by chance.

**Figure0 4.**
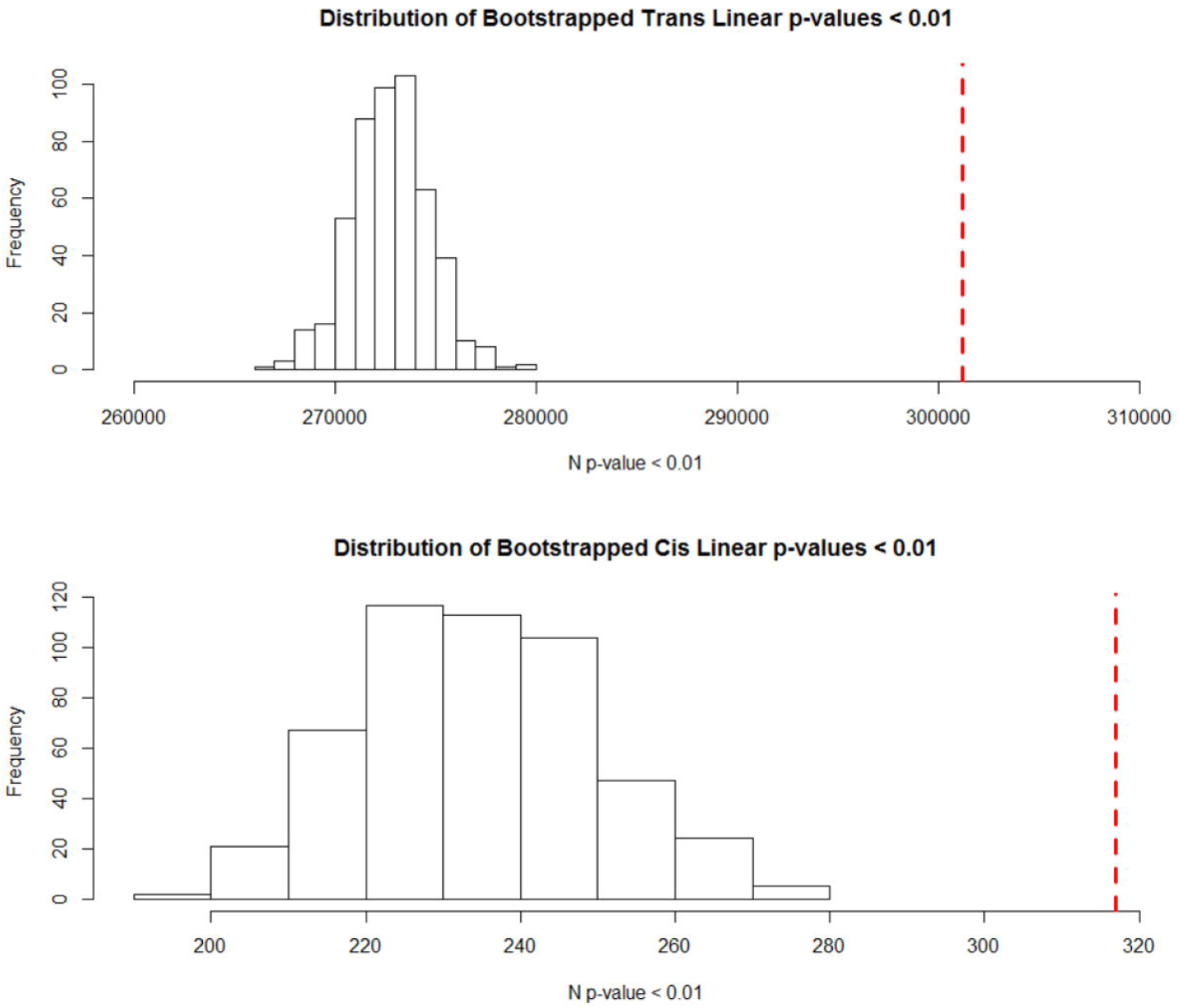
Histograms of the number of p-values below 0.01 in our 500 bootstrapped linear trans and cis-eQTLs analysis. The red dotted line represents the observed values. The likelihood of observing such extreme values by chance is essentially 0 in both cases, if we model the likelihood based of the normal distribution.

**Table 2.** Probability of observing a lower p-value than the lowest, 10^th^ lowest p-value and 100^th^ lowest p-values in our bootstrapping. In general, in relation to our random eQTLs the empirical data was in the lower end of the bootstrapping, except in the linear cis analysis, but not very significantly. It is interesting to note that by the 100^th^ p-value all the analysis outperform random data. This indicates that we do have true, but perhaps weak effects.

| Model | Min P-value | 10<sup>th</sup> P-value | 100<sup>th</sup> P-value |
| --- | --- | --- | --- |
| Anova Cis | 0.13 | 0.126 | 0 |
| **Anova Trans** | 0.044 | 0.062 | 0 |
| **Linear Cis** | 0.984 | 0.688 | 0 |
| **Linear Trans** | 0.148 | 0.024 | 0.018 |

#### Pathway enrichment analysis

As our genes were pre-selected, there is no a-priori reason to perform enrichment analysis. In particular, there is no particular meaning in finding that the cis-eQTLs are enriched for some pathway. The cis-eQTLs are simply tests of correlation between local genomic context and expression, and significance denotes the identification of possible genetic expression regulatory mechanisms, not underlying pathways. In contrast, for the trans-QTL, there are meaningful hypothesis we could state pathways of genes affected by trans-eQTLs. Why would a gene have significant association to a distal genetic element? Previous studies have looked at the local context of trans-eQTLs (26, 27), however we were not able to find any overrepresentation of our pathways in our data (see table S2). Thus, we hypothesized that genes that interact with genomic context and/or are regulate expression would be enriched in the low p-value group in comparison to the overall genes used in genes targeted by trans-eQTLs. This includes genes that directly interact with genomic context, such as DNA binding genes, and regulatory genes, such as transcription factors and positive or negative expression regulators. To test our hypothesis, we selected the top 28147 SNP-gene pairs from our linear trans-eQTL analysis. This represented our observed surplus of low p-values found when comparing with a uniform p-value distribution for eQTLs with a p-value < 0.01, motivated by our results in from the bootstrapping (figure 2). Traditionally, one might test our hypothesis using a pathway enrichment tool, but given that the eQTL data had a special structure, including repeated entries of the same genes from a smaller background set and a large overall number of genes, it was not suitable for typical methods. Instead, we used a more targeted approach, selecting specific categories we believed tested our hypothesis. In table 3, you can see the result for the enrichment of our top linear trans-eQTL genes compared to the initial background set, based on the exact Fisher test, with selected gene ontologies and categories. The results from the enrichment are quite striking, as we get very significant enrichment for DNA binding genes, transcription factors and DNA binding transcription factor activity. All these categories fit our hypothesis, as they engage directly with distal genomic context. We also tested for nucleus genes, as we expect genes that are active in the nucleus to be more likely to interact with genomic context. Furthermore, we tested for general expression regulation, with the positive and negative expression regulation categories. Intriguingly, positive regulation was slightly depleted or unchanged, while negative expression regulatory genes was the most enriched category. Finally, we included membrane genes, as a control category that includes a large number of genes, as we do not believe they have a reason to be enriched which they are not in comparison to the set of genes we used in the eQTL analysis. As a control of the enrichment, we also compared with all expressed genes in our samples, beyond our selected eQTL analysis set. This aids in the interpretation, and acts as a safeguard, as if there was high divergence in the two comparisons the results might just be an artefact of our methodology. We see similar results comparing with all genes, and due to the large number of genes in both the expressed set and the trans-eQTLs, we get very significant p-values. When comparing to the expressed gene-set, we do see slight enrichment for membrane genes, but it should be noted that the baseline ratio between our test set and expressed set for membrane genes is already 1.05 (see table S1), and small effect are significant with the large number of genes included.

**Table 3.** Enrichment analysis based on the linear trans-eQTLs with p-value < 0.01, based on the Fisher exact test. The enrichment was compared with the original input set of 1425 genes, and to the set of expressed genes (N=13202) in our muscle samples for additional comparison.

| Category | N hits in<br>P<0.01 | Odds Ratio | P-value | Odds ratio<br>expressed genes | P-value<br>expressed genes |
| --- | --- | --- | --- | --- | --- |
| *Transcription*<br>*Factor* | 3145 | 2.27 | $1.0 \times 10^{-13}$ | 1.40 | $< 2.2 \times 10^{-16}$ |
| *DNA binding* | 3394 | 2.20 | $8.9 \times 10^{-14}$ | 1.73 | $< 2.2 \times 10^{-16}$ |
| *DNA-binding*<br>*transcription*<br>*factor activity* | 2721 | 3.36 | $< 2.2 \times 10^{-16}$ | 2.36 | $< 2.2 \times 10^{-16}$ |
| *Positive*<br>*regulation of*<br>*expression* | 346 | 0.67 | 0.07 | 0.67 | $8.9 \times 10^{-6}$ |
| *Negative*<br>*regulation of*<br>*expression* | 1887 | 4.34 | $< 2.2 \times 10^{-16}$ | 5.39 | $< 2.2 \times 10^{-16}$ |
| *Nucleus gene* | 8811 | 1.33 | $2.4 \times 10^{-6}$ | 1.18 | $1.8 \times 10^{-12}$ |
| *Membrane gene* | 7707 | 1.06 | 0.30 | 1.15 | $1.8 \times 10^{-8}$ |

## Discussion

In this study, we applied Matrix eQTL to a set of genes previously identified as having potential relations to FCR. We have presented that top results of both cis and trans-eQTLs based on both a linear association and a factor based analysis (Anova). There have been several muscle eQTL studies in pig before (7, 10, 22, 42–46). However, direct comparison of results is quite challenging, for several reasons. None of the other studies where applied to FCR, as the genes and SNPs selected were generally selected based on the phenotype of interest, this limits the overlap. Furthermore, due to the statistical challenges, many divergent strategies were employed, for example using a pre-GWAS(43), picking a limited set of pathway specific genes(10) or using a limited set of microsatellites(46). Some studies also included heritability analysis (7, 22). The studies above include both crossed, purebred and F2 half-sib pig populations. Given all these factors, and the novelty of FCR in an eQTL context, we cannot compare our study very specifically to others, and one should view our study as a pilot study for FCR eQTLs. We do however present novel strategies in an eQTL context, which show promising results, and could be generally applicable to other eQTL studies.

We have included two pure breeds in our analysis, Duroc and Landrace, which in of itself is an unusual choice. Many studies published have inbred lines, but it has been suggested that it would be advantageous to do eQTL analysis on a natural genetically varying population (28), such as two separate breeds. For the input SNPs, we made several choices for maximizing the number of relevant SNPs to include. First, we selected a quite high cutoff of 0.3 MAF. This allows us to have high enough variation at each included SNP, given our low sample size. It has also been shown, that in chip-based data such as ours, the overall structure in the data is robust to different MAF cutoffs(47), thus this should not impart any biases into the results. Finally, we grouped SNPs in high LD (R^2^ >0.9) into blocks and used tagging variants to represent blocks. This allowed us to reduce the space further, removing redundant genetic information, thus relaxing our multiple testing thresholds. In regards to our cis-eQTL distance, we chose a 1mb window, which is on the lower end for pig studies (22). Given our relatively low samples size we wanted to keep the cis analysis as conservative as possible.

In regards to individual eQTLs, one should be careful with over interpreting the results, but instead view the eQTLs as candidates for further study. Based on a qualitative analysis, we do find several interesting genes among our top eQTL candidates. IFI6, a gene implicated in apoptosis regulation through mitochondrial pathways(48), has been previously related to meat and carcass quality(49). PRPF39, a pre-mRNA processing gene, has previously been related to a trans-eQTL in Durocs with divergent fatness(46). DNAJB1, a heatshock gene, was found to be downregulated in lean pigs (50). TMEM222, a transmembrane protein, was found to be differentially expressed between tissues and genotypes between Korean native pigs and Yorkshires(51). CSRNP1, the Cysteine And Serine Rich Nuclear Protein 1 gene, was found to be a metabolic response gene in relation to feed intake in Durocs. INTS7, an RNA processing gene, was associated with a SNP significant for meat quality in Chinese pigs(52). The ACOX3 gene, a fatty acid metabolism gene, had previous cis-eQTLs identified associated with it (10). The PARK7 gene, a gene that codes for a protein that protects from oxidative stress(53), does not appear in a pig related context in the literature, but it is found in a known FCR QTL region, as well as the mitochondrial fission factor (MFF). CPT1B, the arnitine palmitoyl transferase 1B gene, was differentially expressed in large whites versus an indigenous high-fat breed(54). The Potassium Calcium-Activated Channel Subfamily M Alpha 1 gene, KCNMA1, had been previously found to diverge in expression between Large White and Basque pigs(55). Metaxin (MTX2), a mitochondrial gene, was a candidate gene for red blood cell count in a Duroc x Erhualian population based on a nearby genome wide significant SNP (56). Synaptotagmin 12 (SYT12) was found in the area with which explained the largest variance in piglets porn in 3520 Durocs(57). Myosin XIX (MYO19) was a candidate gene for eating behavior traits due to a nearby significant SNP in the same Duroc population our pigs come from(58). The Uncharacterized Protein C7orf50 had a previous cis-eQTLs identified in a behavioral context in humans(59). While this might seem like a mixed group of results, the main takeaway is that each of the genes mentioned above have appeared in previous contexts that demonstrate genetic regulation and association with traits under selection in commercial pigs, thus giving qualitative evidence that increases the likelihood of our eQTLs being true positives.

The final and perhaps most interesting result in our analysis stems from the enrichment analysis in the linear trans-eQTL analysis. We had initially hypothesized that we would find enrichment for genes that interact with genomic context and highly interacting genes. The findings, and their significance level, show a strong overrepresentation of DNA-binding genes, DNA-binding with transcription factor activity genes and transcription factors. These results have a quite straightforward interpretation - genes that interact on a genomic level have a higher chance of having trans-eQTL activity. This can be both mediated through direct interactions, but also through indirect effects, such as transcription factor acting on each other, thus mediating their own genetic effect to other genes. The more intriguing result is the contrast between negative and positive gene regulation. It may be that there is a gene regulation mechanism correlated to trans-eQTLs that explains why we have such a high enrichment of negative regulation, but given our sample size and study power, it is difficult to assess individual genes, and thus properly grasp specific interpretation of these results. The general pathway results are very statistically significant, indicating that these effects should be observable in other eQTL studies. In general, given the complexity of gene expression regulation, further specific study is needed before we have a proper understanding of the contrast between negative and positive expression, and the rest of the enrichment results. Based on our analysis, we propose that these enrichments could be general effects, and thus can assist us in the validation of true trans-eQTLs.

Essentially, identification of such biologically relevant effects can be used as an extra layer of evidence for true positive trans-eQTLs. If we view these results in an animal breeding and selection context, it shows that there may be a merit to weigh the importance of genetic variation based on the gene and pathway context, and weighted methods have been shown to improve the accuracy of breeding value estimates.

## Conclusion

The analysis of eQTLs is a statistically challenging but powerful method for the functional analysis of genetic variation. Here, using pigs from two breeds and different selection goals, we were able show that is possible to identify signal of true positive cis and trans-eQTLs by applying meaningful filtering measures and bootstrapping to show significance. Furthermore, we were able to show highly significant enrichment of regulatory and DNA binding genes in trans-eQTLs, acting as strong evidence for the validity of our trans-eQTLs, and as evidence for a general hypothesis of the nature of trans-eQTLs.

## Acknowledgements

The Authors thank the SEGES – Pig Research Centre (VSP) Denmark and Danish Crown for access to samples and phenotype datasets.

## Contributions

HK conceived and designed this “FeedOMICS” project, obtained funding as the main applicant Independent Research Fund Denmark (DFF). VC and HK designed the blood sampling experiments, phenotype data collection, and biostatistical/bioinformatics analyses. VC carried out bioinformatic data analyses. All authors collaborated in the interpretation of results, discussion, and write up of the manuscript. All authors have read, reviewed, and approved the final manuscript.

