## Supplementary File for "Genetic variations (eQTLs) in muscle transcriptome and mitochondrial genes, and trans-eQTL molecular pathways in feed efficiency from Danish breeding pigs"

| **Category** | **Odds Ratio** | **P-value** |
| --- | --- | --- |
| *Transcription Factor* | 0.62 | 8.4x10^-5^ |
| *DNA binding* | 0.83 | 0.14 |
| *DNA-binding transcription factor activity* | 0.72 | 0.042 |
| *Positive regulation of expression* | 0.98 | 1 |
| *Negative regulation of expression* | 1.33 | 0.20 |
| *Nucleus gene* | 0.97 | 0.70 |
| *Membrane gene* | 1.04 | 0.55 |

*Table S1 – Fisher exact test for selected pathway enrichment of our genes included in the eQTL analysis and our set of total expressed genes.*
