## Supplementary File 2 for "Genetic variations (eQTLs) in muscle transcriptome and mitochondrial genes, and trans-eQTL molecular pathways in feed efficiency from Danish breeding pigs"

| **Category** | **Odds Ratio** | **P-value** |
| --- | --- | --- |
| *Transcription Factor* | 0.83 | 6.1x10^-6^ |
| *DNA binding* | 0.86 | 5.3x10^-4^ |
| *DNA-binding transcription factor activity* | 1.05 | 0.37 |
| *Positive regulation of expression* | 1.11 | 0.20 |
| *Negative regulation of expression* | 1.20 | 0.056 |
| *Nucleus gene* | 0.70 | <2.2x10^-16^ |
| *Membrane gene* | 1.02 | 0.31 |

*Table S2 – Fisher exact test for selected pathway enrichment for the nearest genes to our trans-eQTLs with P < 0.01 compared with our set of total expressed genes.*
